# Mass Spectrometry Reveals that Oxysterols are Secreted from Non-Alcoholic Fatty Liver Disease Induced Organoids

**DOI:** 10.1101/2023.02.22.529551

**Authors:** Kristina Sæterdal Kømurcu, Ingrid Wilhelmsen, James L Thorne, Stefan Johannes Karl Krauss, Steven Ray Haakon Wilson, Aleksandra Aizenshtadt, Hanne Røberg-Larsen

## Abstract

Oxysterols are potential biomarkers for liver metabolism that are altered under disease conditions such as non-alcoholic fatty liver disease (NAFLD). We here apply sterolomics to organoids used for disease modeling of NAFLD. Using liquid chromatography-mass spectrometry with on-line sample clean-up and enrichment, we establish that liver organoids produce and secrete oxysterols. We find elevated levels of 26-hydroxycholesterol, an LXR agonist and the first oxysterol in the acidic bile acid synthesis, in medium from steatotic liver organoids compared to untreated organoids. Other upregulated sterols in medium from steatotic liver organoids are dihydroxycholesterols, such as 7α,26–dihydroxycholesterol, and 7α,25-dihydroxycholesterol. Through 26-hydroxycholesterol exposure to human stem cell-derived hepatic stellate cells, we observe a trend of expressional downregulation of the pro-inflammatory cytokine CCL2, suggesting a protective role of 26-hydroxycholesterol during early-phased NAFLD disease development. Our findings support the possibility of oxysterols serving as NAFLD indicators, demonstrating the usefulness of combining organoids and mass spectrometry for disease modeling and biomarker studies.

**Graphical abstract:** 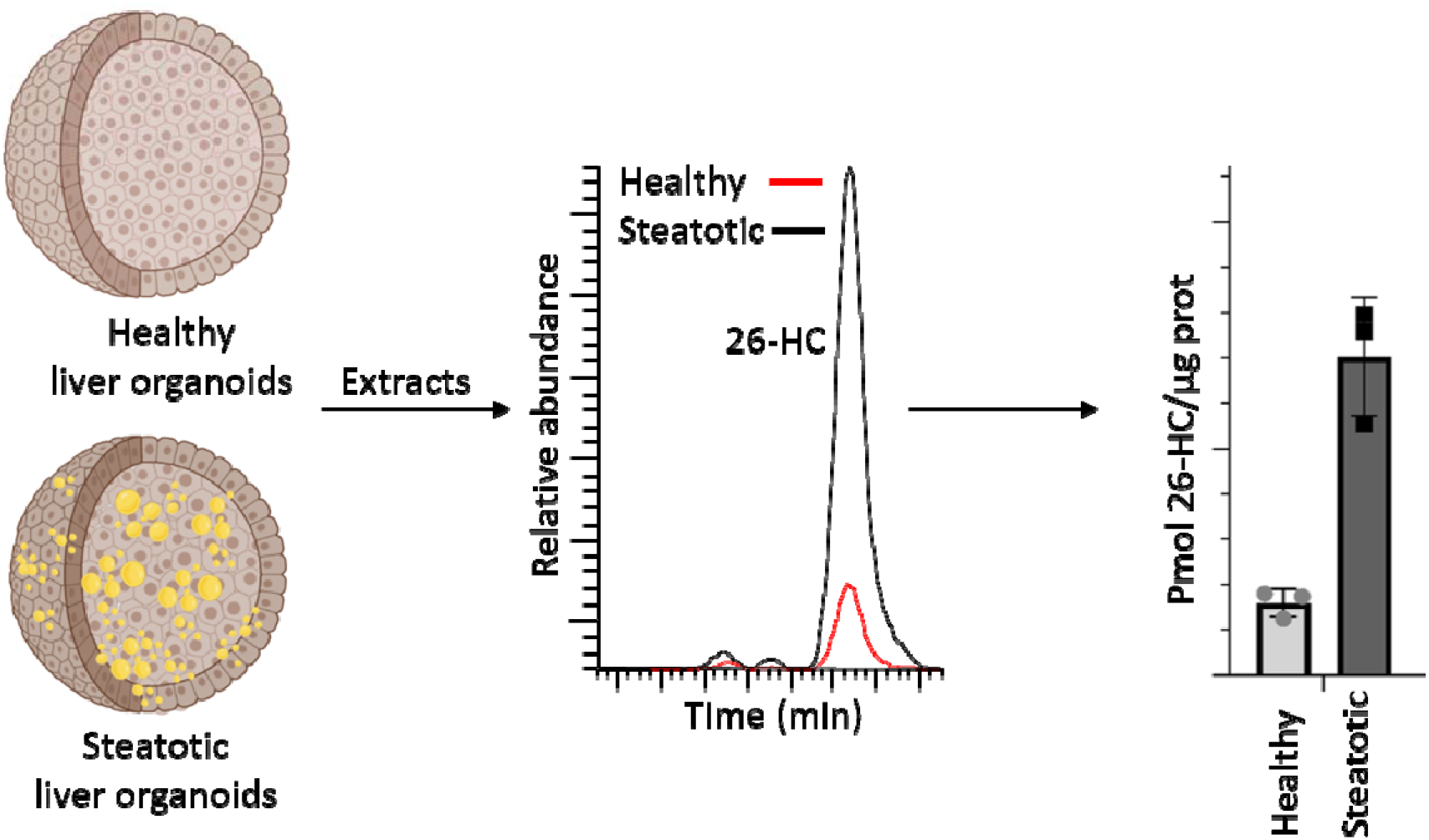

## INTRODUCTION

Non-alcoholic fatty liver disease (NAFLD) is a heterogeneous disease related to the accumulation of high levels of lipids in the liver [1, 2]. Studies estimate the pooled global prevalence of all forms of NAFLD to be 30% [3]. NAFLD is closely interlinked with obesity, but studies also show that diabetes increases the risk of developing more severe stages of NAFLD [2]. NAFLD is divided into stages according to disease progression (*Figure 1*), starting with the benign steatotic stage where the liver has begun to build up lipids before evolving to non-alcoholic steatohepatitis (NASH), where inflammation occurs. As a consequence, tissue scarring (fibrosis) occurs until it reaches cirrhosis, the stage where liver function is significantly severed. Disease progression can cause liver cancer (hepatocellular carcinoma (HCC)) and liver failure ultimately requiring liver transplantation [1].

**Figure 1.**
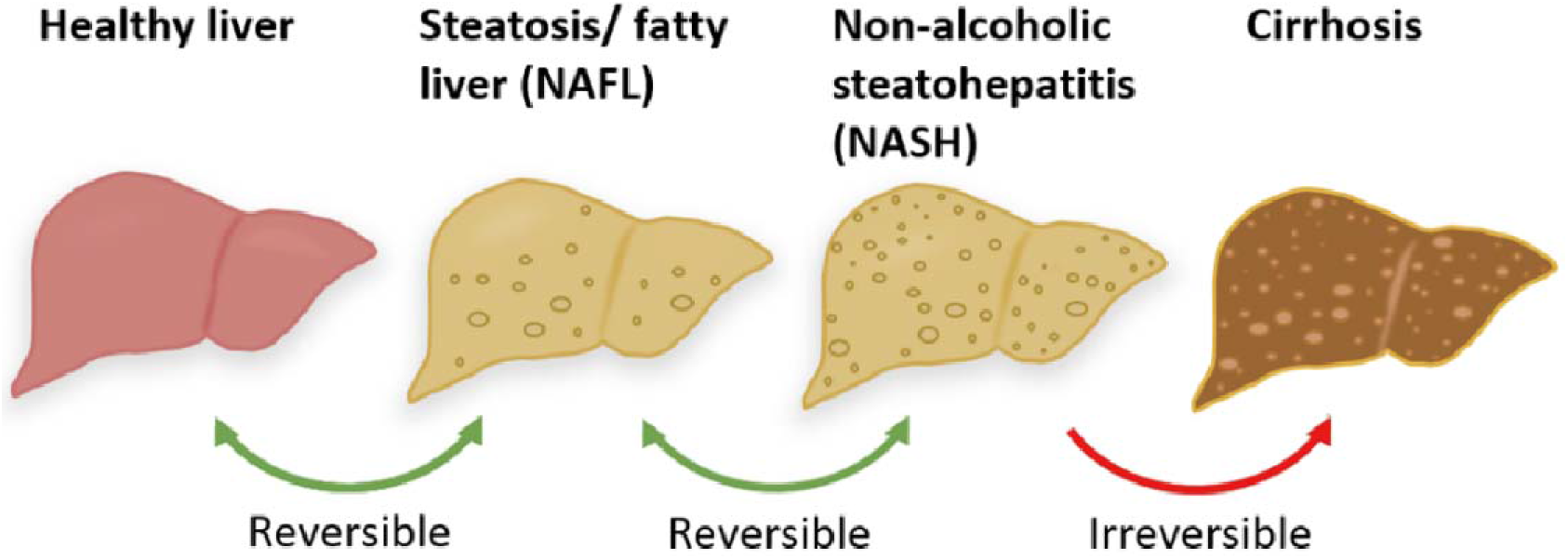
The stages of NAFLD progression. In the first stage, steatosis, the liver accumulates higher levels of fat (fatty liver). The next stage is NASH, where the degree of fibrosis (tissue scarring) is increasing. When the tissue scarring is severe, the liver function is altered (cirrhosis) and liver transplantation is often needed.

Today, the most specific and common test to diagnose and stage NAFLD is a liver biopsy. The use of ultrasound-based approaches or non-invasive diagnostic methods based on biomarkers has pitfalls, such as low sensitivity in early stages leading to an inability to assess the severity of the disease [4]. New biomarkers to increase understanding of disease onset, progression, and diagnosis are needed.

Due to differences between animal models and human physiology - rodent models don’t fully reproduce human liver metabolism [5], both human liver spheroids and liver organoids, are increasingly used for NAFLD modeling and biomarker discovery [6]. While liver spheroids can be described as 3D clusters of differentiated cells without recapitulation of elements of tissue-like structure, liver organoids are developed from human pluripotent stem cells (PSC)[7]. Additionally, stem-cell-derived 2D monocultures of key effector cells can be used for modeling NAFLD in an even more simplified system.

Cholesterol derivates have received attention as potential biomarkers of NAFLD [8–10]. Oxysterols are oxidized forms of cholesterol (Figure 2) and are involved in a plethora of biochemical processes and diseases [11]. In the context of NAFLD, oxysterols are regulators of the liver x-receptor (LXR) [12], a key modulator in NAFLD [13]. Chronic LXR activation contributes to the development of steatosis, where dysregulation of oxysterols metabolism may affect the severity of NAFLD toward NASH [14]. Ikegami et al. found elevated levels of 25-hydroxycholesterol (HC) and 26-HC (also known as 27-HC [15]) in serum samples from NAFLD patients [9]. Raselli et al. found elevated levels of 24- and 7-hydroxylated oxysterols in liver biopsy samples of patients with NASH [10]. Studying murine models of NASH Raselli et al found elevated levels of 24-HC and 7α-hydroxycholeste-4-en-3-one, but these findings differ from studies of the human liver. Hence, while oxysterols are implicated in NAFLD progression and may serve as biomarkers, their exact role in disease initiation and progression in humans remains unclear. Specifically, the effect of oxysterols on liver non-parenchymal cells, essential for the progression of NAFLD, including hepatic stellate cells (HSCs), is unexplored. Hence, studying oxysterols in organoid models of NAFLD may add to our understanding of this metabolites group.

**Figure 2.**
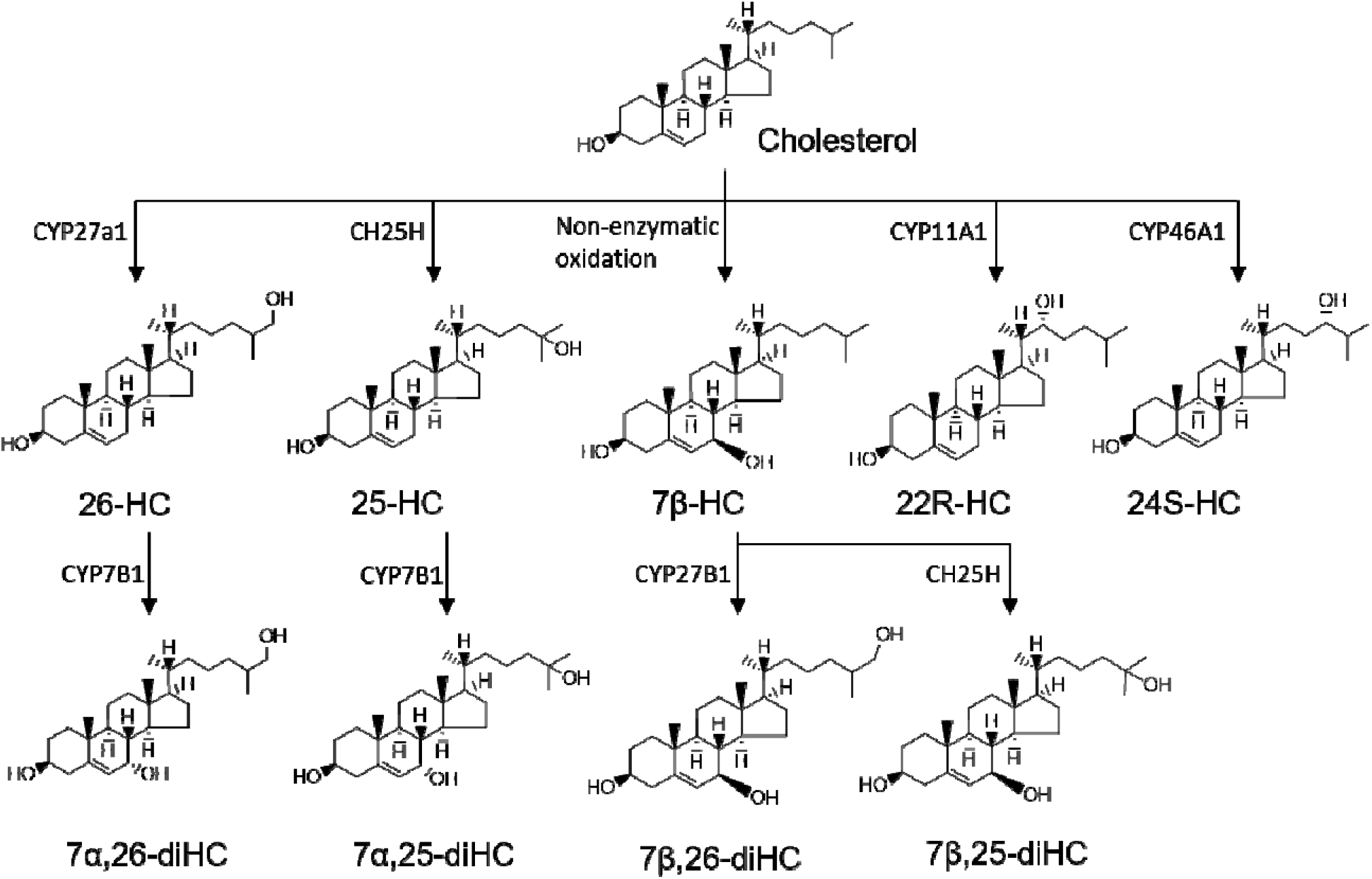
Oxysterol formation. Cholesterol is oxidized to form oxysterols, via cytochrome P450 (CYP) enzymes.

Oxysterols are a highly challenging group of analytes to measure and elucidate, even when applying high-performance liquid chromatography-mass spectrometry (LC-MS). Many oxysterols are isomers, requiring high chromatographic resolution before MS. In addition, oxysterols are difficult to charge, resulting in poor sensitivity when applying conventional electrospray ionization before MS detection. Therefore, sensitive LC-MS requires oxysterols to be derivatized by introducing chargeable groups. To the authors’ knowledge, oxysterols have hitherto not been analysed in human liver organoids and *in vitro* models of NAFLD.

We here introduce mass spectrometric analysis of oxysterols on human liver spheroids and organoids with the goal of assessing correlations between levels of specific oxysterols and early stages of NAFLD (steatosis).

## EXPERIMENTAL

### Generation of human primary hepatic spheroids (PHH)

Cryopreserved primary human hepatocytes (PHH) (Lonza, catalogue no.HUCPG, lot HUM180201A (PHH_1) and Gibco, catalogue no. HMCPMS, lot HU8287 (PHH_2)) were thawed in the Hepatocytes thaw media (Gibco, catalogue no. CM7500) according to the manufacturer’s protocol. Uniform PHH spheroids were created by aggregation in-house-made agarose microwells - a format in which PHH showed stable functionality over at least 7 days as described before[7]. Briefly, cells were plated into microwells at the concentration of 1000 viable cells per microwell and were centrifuged at 100 g for 2 min. For the first 3 days PHH was cultured in the Williams E medium (Thermo Fisher Scientific, catalogue no. A1217601) supplemented with 7 % FBS (Thermo Fisher Scientific, catalogue no. 41400045), 2 mM L-glutamine (Thermo Fisher Scientific, catalogue no. 35050038), 10 μg/ml insulin, 5.5 μg/ml transferrin, 6.7 ng/ml sodium selenite (Thermo Fisher Scientific, catalogue no. 41400045) and 0.1 μM dexamethasone (Sigma Aldrich, catalogue no. D4902). From day 4 onwards, the FBS concentration was gradually decreased to 1 % (v/v).

### Generation of human induced PSC/human embryonic stem cell-derived human-like hepatocytes (HLC)

Human induced pluripotent stem cells (hiPSC)(HLC_1: WTSIi013-A; HLC_3: WTC-11, Coriell Institute for Medical Research) and human embryonic stem cells (hESC) (HLC_2: H1) were cultured in E8 media (Thermo Fisher Scientific, catalogue no. A1517001) on plates coated with 0.1 % (v/v) Geltrex (Thermo Fisher Scientific, catalogue no. A1413201) in a humidified 37 °C, 5 % CO_2_ incubator. Quality control, performed after the thawing of cells, including flow cytometry, real-time polymerase chain reaction, immunofluorescent imaging for pluripotency markers, and karyotyping.

HLC was generated using an protocol established in our laboratory [16]. Briefly, iPSC was differentiated toward definitive endoderm with the subsequent treatment with 10 ng/mL FGF2 (Peprotech, catalogue no. 100-18B) and 20 ng/mL BMP4 (Peprotech, catalogue no. 120-05) in IMDM/F12 medium supplemented with 1% (v/v) N-2 (Thermo Fisher Scientific, catalogue no. 17502-048), 1% (v/v) B-27 minus vitamin A (Thermo Fisher Scientific, catalogue no. 12587010) and 1% (v/v) lipid concentrate, then with 5 μM A8301 (Stem Cell Technologies, catalogue no.72022), 20 ng/mL HGF (Peprotech, catalogue no. 100-39H), 20 ng/mL BMP4, 1% (v/v) B-27 with vitamin A for 3 more days and with 25 ng/mL HGF, 1% (v/v) DMSO for another 5 days. Obtained hepatic progenitors (day 12) were detached and aggregated in the agarose U bottom microwells in the presence of 25 ng/mL HGF, 0.1 μM Dexamethasone, 0.5% (v/v) ITS, 0.1% (v/v) lipids concentrate, 100 μM Ascorbic acid-2 phosphate (AAP), 1% (v/v) B-27 (without vitamin A) and 1% (v/v) N-2. After the formation of spheroids at day 13, media was replaced with William’s E media, supplemented with 5% (v/v) FBS, 20 ng/ml HGF and 10 ng/ml Oncostatin M (Peprotech, catalogue no. 300-10), 1% (v/v) ITS, 100 μM AAP, 0.1 μM dexamethasone. Organoids were cultured in microwells in William’s E media, supplemented with 1% (v/v) ITS, 0.1 μM Dexamethasone, 20 ng/ml Oncostatin M and 1% (v/v) MEM Non-Essential Amino Acids Solution (Thermo Fisher Scientific, catalogue no. 11140050), for another 10 days.

### Steatosis induction

To induce hepatic steatosis, PHH or HLC were incubated for 48 h in the serum-free William’s E media, supplemented with 10 mM fructose, and saturated (palmitic acid, 250 μM) and mono-unsaturated (oleic acid, 250 μM) free fatty acids in ratio 1:1, termed below as steatotic conditions [7]. Incubation was performed in ultra-low attachment 24 wells-plate, with 20-30 organoids per well filled with 500 μl of control or steatotic media.

### Differentiation of human iPSC-/ hESC-derived hepatic stellate cells

2D cultures of human stem cell-derived hepatic stellate cells (sc-HSCs) were generated using modifications of previously published protocols [17, 18]. Human PSCs were seeded as single cells in a 1:4 ratio on plates coated with 1 % (v/v) GelTrex matrix (Thermo Fisher Scientific) and pretreated with 10 μM Rock inhibitor (STEMCELL Technologies). The sc-HSCs differentiation protocol was initiated by the incubation with DMEM/F12 GlutaMax™ medium (Gibco, catalogue no. 31331-093) supplemented with 1 % (v/v) MEM non-essential amino acids (Gibco, catalogue no. 11140050),1 % (v/v) B-27™ supplement (Gibco, catalogue no. 17504044), 100 ng/mL activin A (Peprotech, catalogue no. 120-14P), 3 μM CHIR99021 (Tocris Bioscience, catalogue no. 4423) and 20 ng/mL BMP-4 (Peprotech, catalogue no. 120-05) for 24 hours on day 0 of the differentiation protocol. Mesodermal progenitors obtained in this stage were incubated in DMEM/F12 GlutaMax™ medium supplemented with 1 % (v/v) MEM non-essential amino acids, 1 % (v/v) B-27™ supplement, 1 % (v/v) Insulin-Transferrin-Selenium (Gibco, catalogue no. 41400045), 2.5 μM Dexamethasone (Sigma-Aldrich, catalogue no. D4902-100mg) and 100 μM 2-Phospho-L-ascorbic acid trisodium salt (Sigma-Aldrich, catalogue no. 49752) including 20 ng/mL BMP-4, and, to induce a liver mesenchymal submesothelial phenotype. Next 20 ng/mL FGF-1 (Peprotech, catalogue no. 100-17A) and 20 ng/mL FGF-3 (R&D Systems, catalogue no. 1206-F3-025) were added on days 3-11. On days 6-12, B-27™ supplement was replaced with 1 % (v/v) Fetal Bovine Serum (Gibco, catalogue no. 26140079) and 100 μM Palmitic Acid (Sigma-Aldrich, catalogue no. P5585) and 5 μM Retinol (Sigma-Aldrich, catalogue no. R7632) was added to the media to induce a fetal HSC-like phenotype.

### sc-HSC activation and treatment with cholesterol and oxysterols 25-HC and 26-HC

After differentiation, sc-HSCs were treated with 100 nM 25-HC, 100 nM 26-HC (both from Sigma-Aldrich, St. Louis, MA, USA), or 50 μM cholesterol (Promega, catalogue no. J322A) both with and without 25 ng/mL TGF-β1 (Abcam, catalogue no. ab50036) for 48 hours. Samples were then collected after detachment with Trypsin-EDTA solution (Sigma-Aldrich, catalogue no. T4049-100ML) and snap frozen at −80 °C until RNA extraction and real-time quantitative polymerase chain reaction (RT-qPCR) analysis. Biological triplicates were collected for each condition.

### RNA extraction and real-time quantitative polymerase chain reaction

RNA was isolated using RNeasy Micro kit (Qiagen) or TRIzol reagent (Thermo Fisher) according to the manufacturer’s protocol. RNA concentration and purity were determined using NanoDrop ND-1000 spectrophotometer (Thermo Fisher Scientific). cDNA was synthesized using High-Capacity cDNA Reverse Transcription Kit (Thermo Fisher Scientific, catalogue no. 4368814). Gene expression analysis was performed using a TaqMan Universal mix on a TaqMan ViiA7 Real-Time PCR System. Used primers are listed in the Supporting Information table 1. GAPDH was used as endogenous control. The level of expression of genes of interest was quantified by ddCt with to HLC differentiated from the WTSIi028-A iPSC line (HLC_1), and with normalization to control organoids. Data represent two donor PHH samples, and HLC differentiated from 3 cell lines.

### Immunofluorescence and lipids detection

Organoids were fixed in 4 % (w/v) PFA for 30 min on the orbital shaker. Each step in the protocol was followed by 3 washings (each 5 min) in DPBS using an orbital shaking. Permeabilization and blocking were performed by incubation in PBS with 1 % (w/v) BSA (Sigma Aldrich), 0.2 % (v/v) Tritox-X100 (Sigma Aldrich), and 0.5 % (v/v) DMSO at RT for 2 h on the orbital shaker. Staining with primary antibodies was performed for 24 h (at 4 °C) with subsequently 2 h incubation with secondary antibodies (Jackson ImmunoResearch, West Grove, PA) diluted with 1 % (w/v) BSA, 0.1 % (v/v) Tritox-X100 in PBS. Primary antibodies (Ab) used in this study: rabbit polyclonal Ab to human serum albumin (Abcam, catalogue no.ab2406, 1:400), goat polyclonal Ab to PDGFR-β (R&D Systems, catalogue no. AF385), rabbit monoclonal Ab to NCAM1 (Abcam, catalogue no. ab75813), mouse monoclonal Ab to Vimentin (Abcam, catalogue no. ab8978); secondary Ab used in this study: Cy™3 AffiniPure Donkey Anti-Rabbit IgG (H+L) (catalogue no.711-165-152, 1:400), Alexa Fluor® 647 AffiniPure Donkey Anti-Mouse IgG (H+L) (catalogue no.715-605-150, 1:400) (all secondary Ab are from Jackson ImmunoResearch). Nuclear counterstaining was performed with 1 μg/mL Hoechst 33258 (Sigma Aldrich) for 5 min at RT. Neutral lipids were stained using HCS LipidTOX™Red Neutral Lipids (excitation/emission maxima ~577/609 nm, Thermo Fisher Scientific, catalogue no. H34476) or Nile Red (Sigma Aldrich).

Confocal microscopy was performed on a Zeiss 700 laser scanning confocal microscope. Confocal microscope using standard filter sets and laser lines with a 40x oil immersion objective. Images were acquired using Zen software (Zeiss) as Z-stacks with 2 μm spacing between stacks and Dragonfly software with 0.5 μm spacing correspondingly. The confocal images were analysed using Fiji software [19]. Confocal images are displayed as a resulting Z-stack of half of the spheroid.

### Detection of vitamin A in HSCs

Vitamin A storage was indirectly determined by imaging autofluorescence in the stem cell-derived HSC under UV light excitation. Cells were imaged using a 10X magnification objective on an Axioscope fluorescent microscope (Zeiss, Jena Germany).

### LC-MS analysis

#### Chemicals

Methanol (MeOH, Hipersolv), acetonitrile (ACN, LC-MS grade), and sodium hydroxide (NaOH) were from VWR (Radnor, Pa, USA), and type 1 water was from a Millipore integral system (Millipore, Billerica, MA, USA). Girard T reagent, cholesterol oxidase from *Streptomyces sp*., KH2PO4, formic acid (FA, HPLC-grade, 98 %), 22R-HC, 25-HC, and cholesterol-25,26,27-^13^C_3_ were all purchased from Sigma-Aldrich (St. Louis, MA, USA). A stock solution of 50 mM phosphate buffer with 0.03 mg/mL cholesterol oxidase (pH adjusted to 7 with 1 M NaOH) was stored at −80 °C; fresh buffer was thawed before each use. Glacial acetic acid was from Merck (Merck KGaA, Darmstadt, Germany). 7β,25-diHC, 7α,25-diHC, 7α,24S-diHC, 7α,26-diHC, 7β,26-diHC, 24S-HC, 26-HC, 7α,25-diHC-d_6_, 7α,26-diHC-d_6_, 25-HC-d_6_ and 26-HC-d_6_ were purchased from Avanti Polar Lipids (Alabaster, AL, USA). All stock solutions of standards or internal standard (95-100 μM) were prepared in 2-propanol (LC-MS-grade, Rathburn Chemicals Ltd., Walkerburn, Scotland, UK) and stored at −20 °C.

#### Calibration solutions

Working solutions of 1 nM standard mixture A (7β,25-diHC, 7α,25-diHC and 7α,24S-diHC), 1 nM standard mixture B (7α,26-diHC, 7β,26-diHC, 22R-HC, 25-HC. 24S-HC, and 26-HC) and 1.5 nM internal standard mixture (7α,25-diHC-d_6_, 7α,26-diHC-d_6_, 25-HC-d_7_ and 26-HC-d_7_) were diluted from 95-100 μM stock solutions. Calibration *solutions* in the concentration range of 50-500 pM (200 pM internal standard) were made of 100 μL internal standard mixture, 37-370 mL standard mixture (A or B), 100 μL media (for matrix matching) and 88 μL of 1 nM cholesterol-25,26,27-^13^C_3_ (120 pM final concentration, for autoxidation monitoring). Samples were evaporated to dryness and resolved in 20 μL IPA before Girard T-derivatization. New calibration solutions were made for each assay.

#### Liver organoid samples

For each cell line three replicates were collected and again divided into three technical replicates, providing 9 replicates for each cell line. Approximately 90 μL of the organoid medium was mixed with 100 μL internal standard (200 pM) and 88 μL of 1 nM cholesterol-25,26,27-^13^C (120 pM final concentration) for autooxidation monitoring [20]. Samples were evaporated to dryness and redissolved in 20 μL IPA before Girard T-derivatization. All samples were stored at 4 °C and analysed within 3 weeks.

#### Sample preparation – Girard T

Oxysterols were derivatized by the procedure described by Griffith et al [21] with modifications as described by Roberg-Larsen et al [22] (Figure 3). Briefly, 200 μL of aliquoted cholesterol oxidase solution was added to each sample/standard and incubated at 37 °C for one hour. A mixture of 500 μL MeOH, 15 μL glacial acetic acid, and 15 mg Girard T was added to each sample followed by incubation overnight in darkness. Samples were then stored at 4 °C and analysed within 2-3 weeks.

**Figure 3.**
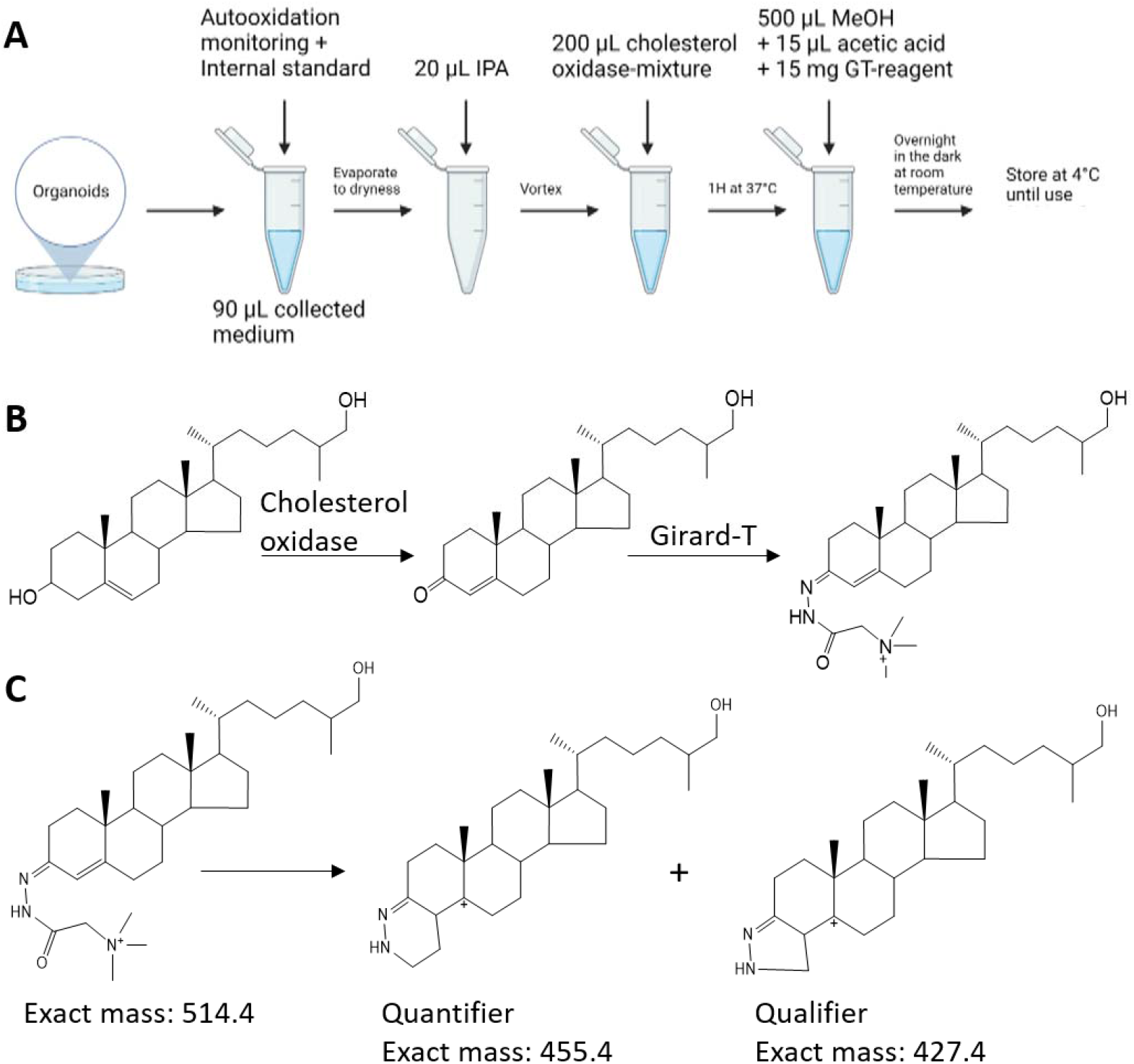
*Sample preparation of oxysterols. Medium is collected from the organoid wells and sample preparation, including Girard T-derivatization, is performed according to the process shown in A. B) The derivatization of 26-HC with Girard T reagent. The OH-group is first oxidized by cholesterol oxidase, before the newly formed carbonyl group reacts with the Girard T reagent, forming a Girard T-derivate. C) The precursor oxysterol-Girard T is fragmented in the MS into the quantifier* (*m/z* = 455.4) and the qualifier (*m/z* = 427.4).

#### Liquid chromatography-mass spectrometry system

A previously established method for the hydroxycholesterols 22R-HC, 25-HC, 24s-HC and 26-HC was altered to also include the five dihydroxycholesterols 7α,25-diHC, 7β,25-diHC, 7α,24S-diHC, 7α,26-diHC and 7β,26-diHC [23]. Mobile phase A was 0.1 % FA in type 1 H_2_O; B was 0.1 FA in MeOH, and C was 0.1 % FA in ACN. Compared to the aforementioned method, the step gradient was extended with a 7 min step of MP composition 62/14/24 (v/v/v, A/B/C) at the start of separation to achieve separation of the diHCs before an increase in MP strength to 56/14/30 (v/v/v, A/B/C). A 2.5 min wash step followed using a MP composition of 50/50 (v/v, B/C). The total method runtime, including on-line sample clean-up and column conditioning, was 21 min. Aliquots of 80 μL of derivatized samples were loaded onto an automated filtration and filter backflush solid phase extraction (AFFL)-system, as described in Solheim et al [23] (Figure 4). Briefly, in the “load”-position, injected samples are filtered on-line before focusing of analytes to an on-line HotSep C18 SPE column (Teknolab, Ski, Norway). Loading is performed at a flow rate of 0.5 μL/min, using mobile phase A, by a Hitachi L-7110 pump (Merck). Non-retained compounds (e.g. excess Girard-T reagent) are sent to waste. When the valve is switched to the “inject” position, the retained compounds are eluted from the SPE and subsequently separated on an ACE SuperPhenyl Hexyl (2.1 mm ID, 150 mm, 2.5 μm d_p_ core-shell particles)-column from Advanced chromatography technologies LTD, Aberdeen, UK (column temperature set to 55 °C) using a Dionex UltiMate 3000 UHPLC-system with a flow rate of 0.7 mL/min (pump 2). A TSQ Vantage triple quadrupole mass spectrometer (Thermo Fisher Scientific, Waltman, MS, USA) operated in selected reaction mode (SRM) was used for MS-analyses; see Supporting Information *Table 1* for MS parameters. Xcalibur software (Thermo) was used for peak integrations.

**Figure 4.**
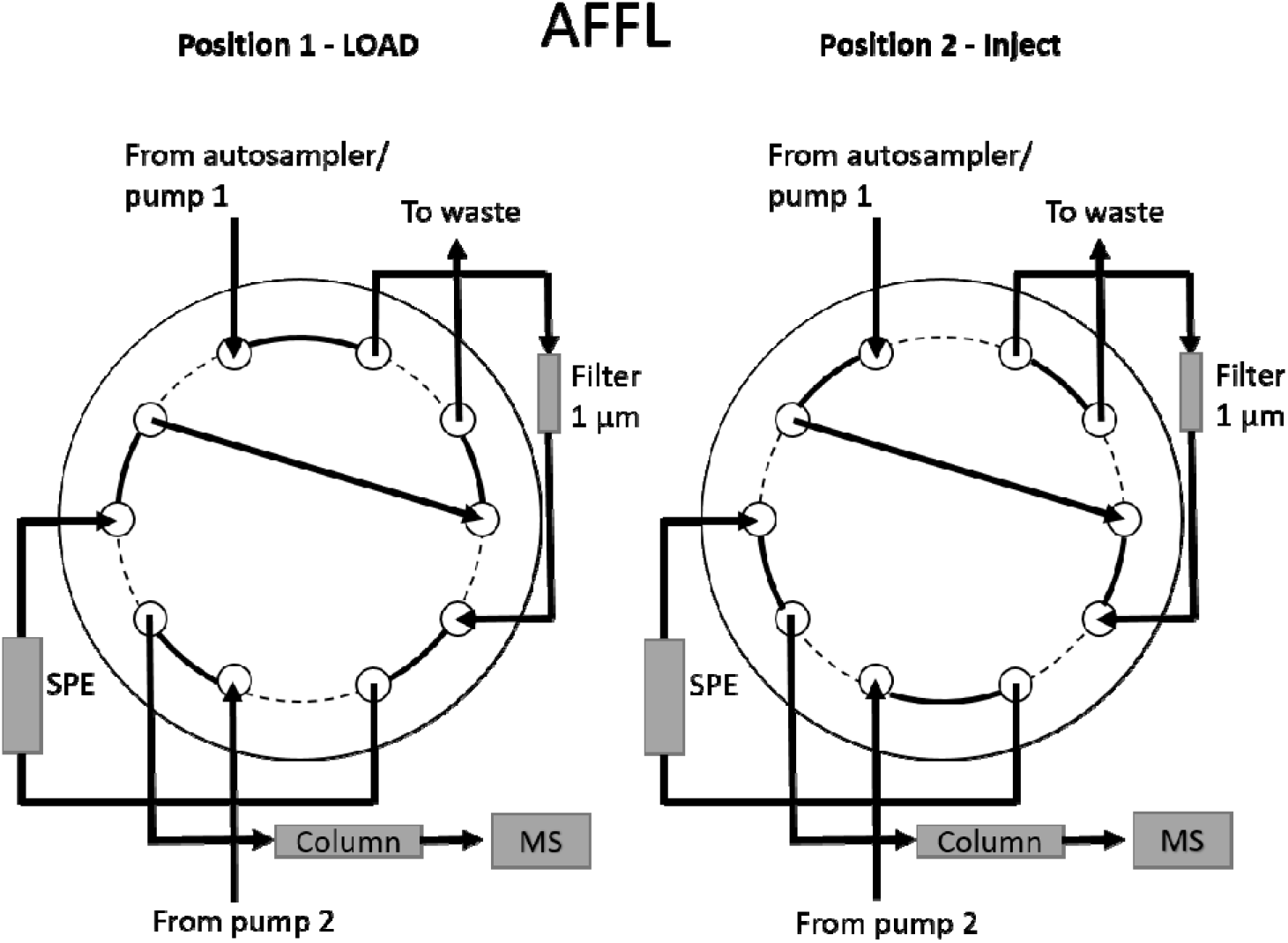
The AFFL-system. Illustration of the automated filtration and filter backflush solid phase extraction system (AFFL), coupled on-line with LC-MS.

#### Method evaluation

The method was evaluated at three different concentration levels for both intra-day and inter-day precision, using spiked organoid medium. For intra-day evaluation, 6 replicates for low concentration (50 pM), medium concentration (200 pM), and high concentration (500 pM) were analysed and relative standard deviation was calculated. One-way ANOVA was used to calculate the inter-day-precision (intermediate precision) based on three replicates of each of the three concentration levels prepared and analysed on 4 different days. The limit of quantification is defined as 10 times the S/N and the limit of detection as 3 times S/N, where the ratios for each peak are defined by XCalibur (set to INCOS Noise). No carry-over was observed.

#### Calculations and statistics

Statistical analyses and graph generation were performed using GraphPad PRISM 9 (GraphPad Software Inc.). For qPRC-analysis, a two-tailed, paired *t*-test was applied for the comparison of two groups. For more than two groups, a one-way ANOVA analysis was applied. For LC-MS results, reported calculations are normalized against the measured protein amount in the corresponding organoid well and reported in pmol per μg protein. When comparing oxysterol levels from steatotic to control organoid samples, a multiple unpaired t-test with Welch correction was used. All data are presented as meanͰ±ͰSD.

## RESULTS AND DISCUSSION

### Chromatographic system

Oxysterols are isomers with highly similar MS fragmentation, requiring chromatographic separations before detection. By modifying our sterolomics platform [23, 24], we were able to separate and quantify four hydroxycholesterol isomers (22R-HC, 24S-HC, 26-HC, and 25-HC) and five dihydroxycholesterol isomers (7α,25-diHC, 7β,25-diHC, 7α,24S-diHC, 7α,26-diHC and 7β,26-diHC) in 21 minutes, including on-line SPE sample clean-up to remove excess derivatization reagent (0.12 M or 20 mg/mL) and cell medium (Figure 5). Using a phenylhexyl stationary phase we observed no peak splitting of *syn/anti* isomers of post-derivatized oxysterols, an issue often encountered with conventional RPLC stationary phases [21]. The method was evaluated in the range of 50-500 pM for each of the nine analytes (Supporting information *Table 2*). All standard solutions were added cell medium used for organoids before sample preparation for matrix matching. Intra-day RSD values were <28 % for “high” concentrations and < 17% for “middle” concentrations for all HC and diHC. For “low” concentrations, all RSDs were below 22 % except 7α,24S-diHC (RSD= 44%). The inter-day RSD values were < 20% for the HCs at all concentrations. For the diHCs inter-day RSDs were higher, ranging from 8% to 20% (7β,25), 12%-27% (7α,25), 17-41 % for 724S, 25-38 % (7β,26) and 31-48 % (7α,26). We experienced two peaks for each of the dihydroxycholesterols in standards, with the latter peak being smaller, but the ratio between the peaks was consistent. The additional peaks co-eluted with the major peak of another dihydroxycholesterols, e.g. the latter peak for 7β,25-diHC co-eluted perfectly with 7β,26-diHC under all chromatographic conditions, implying that it could be due to impurities in the standard as seen earlier [24]. Hence, we found it adequate to make two calibration curves avoiding the impact of impurities from the standards. R^2^ values were > 0.97 for all analytes. The chromatographic resolution ranged from 0.8-1.3 between the isomers. Taken together, the sensitivity, precision, and resolution were considered satisfactory for introducing sterolomics and organoid analysis.

**Figure 5.**
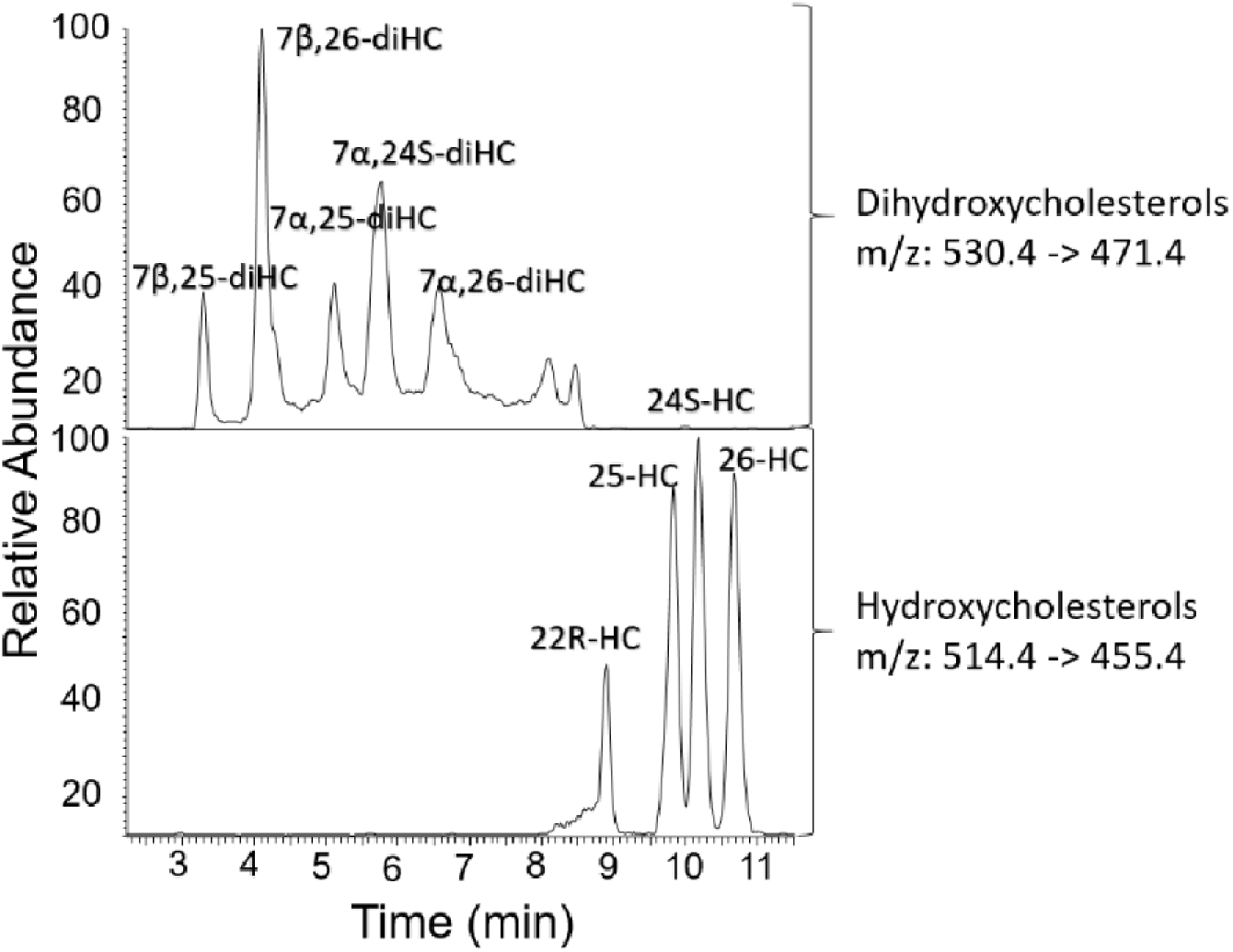
Chromatograms for all nine oxysterols included in this study. Separation is achieved in 11 minutes, and the total run-time for the method including on-line sample cleanup is 21 min. In this figure, a standard solution with a concentration of 500 pM is injected. Cell medium was added at the beginning of the sample preparation for matrix matching. Monitoring of 13C-labelled cholesterol products was also performed; no autoxidation was detected throughout the study.

### Healthy liver spheroids/organoids secrete 26-HC

Applying our method we could detect and quantify 26-HC secreted from 20 liver organoids per sample (*Figure 6*). Medium from both primary human spheroids (two different donors; PHH-1 and PHH-2) and hESC/hiPSC-derived liver organoids (three different cell lines; HLC-1, HLC-2, and HLC-3) contained quantifiable levels of 26-HC (see *Figure 6* for representative chromatograms and *Figure 7* for quantitative data). The presence of 26-HC documents sufficient activity of the enzyme CYP27A1 to render detectable levels of secreted 26-HC. The amounts of 26-HC secreted from control/healthy HLCs are generally less than that secreted from the PHH, which is in accordance with a generally lower level of enzyme activity in HLC than in PHH [25]. In contrast to 26-HC, 25-HC, and 24S-HC were detectable only in some of the samples and at levels that are close to the detection limit. The above-specified dihydroxycholesterols were not detected in any of the samples (Figure 6).

**Figure 6.**
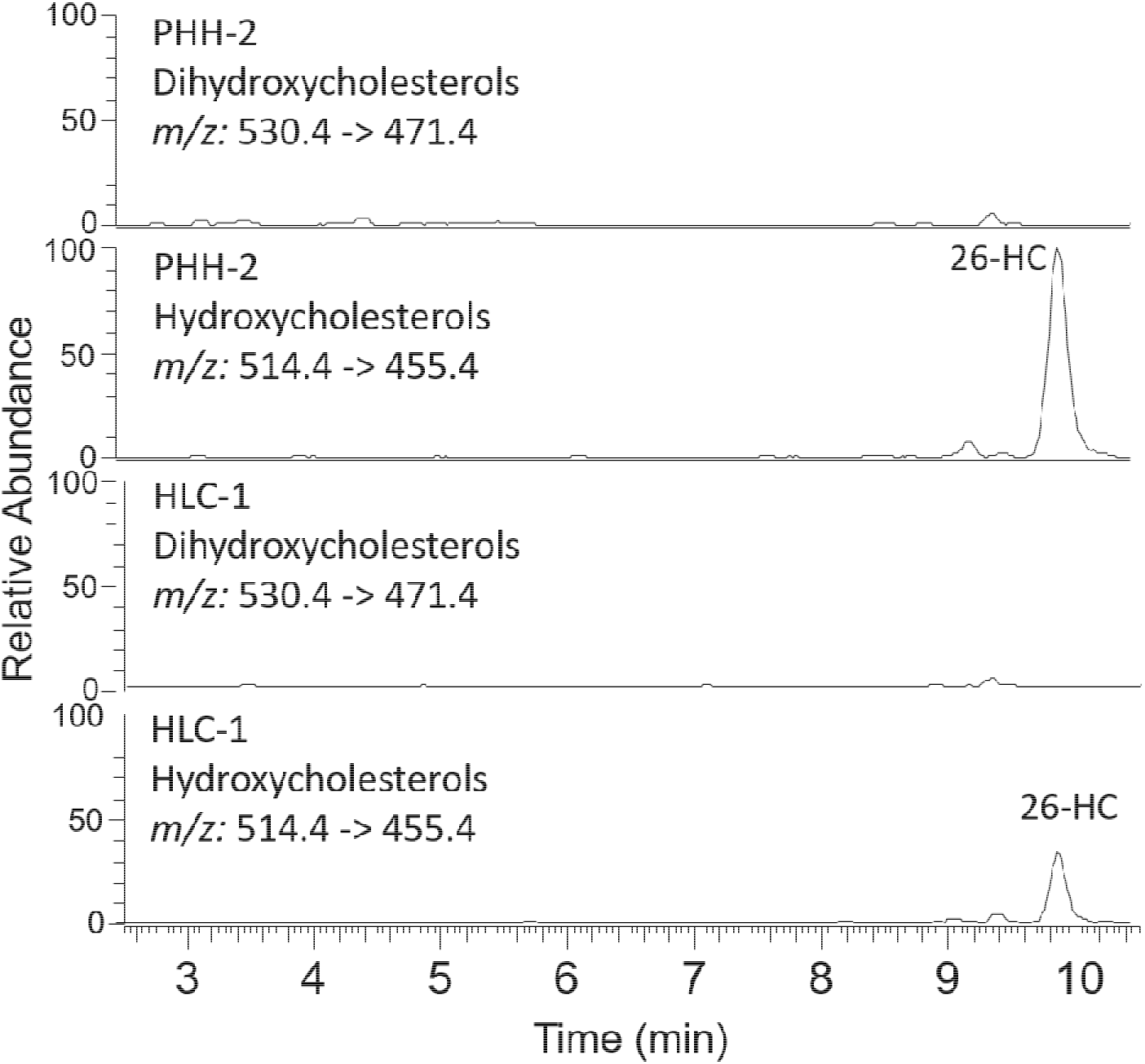
Extracted ion chromatograms for untreated PHH and HLC. For both the m/z- transitions 530.4 → 471.7 and 514.4 → 455.4 is shown, corresponding to the dihydroxycholesterols and hydroxycholesterols, respectively.

**Figure 7.**
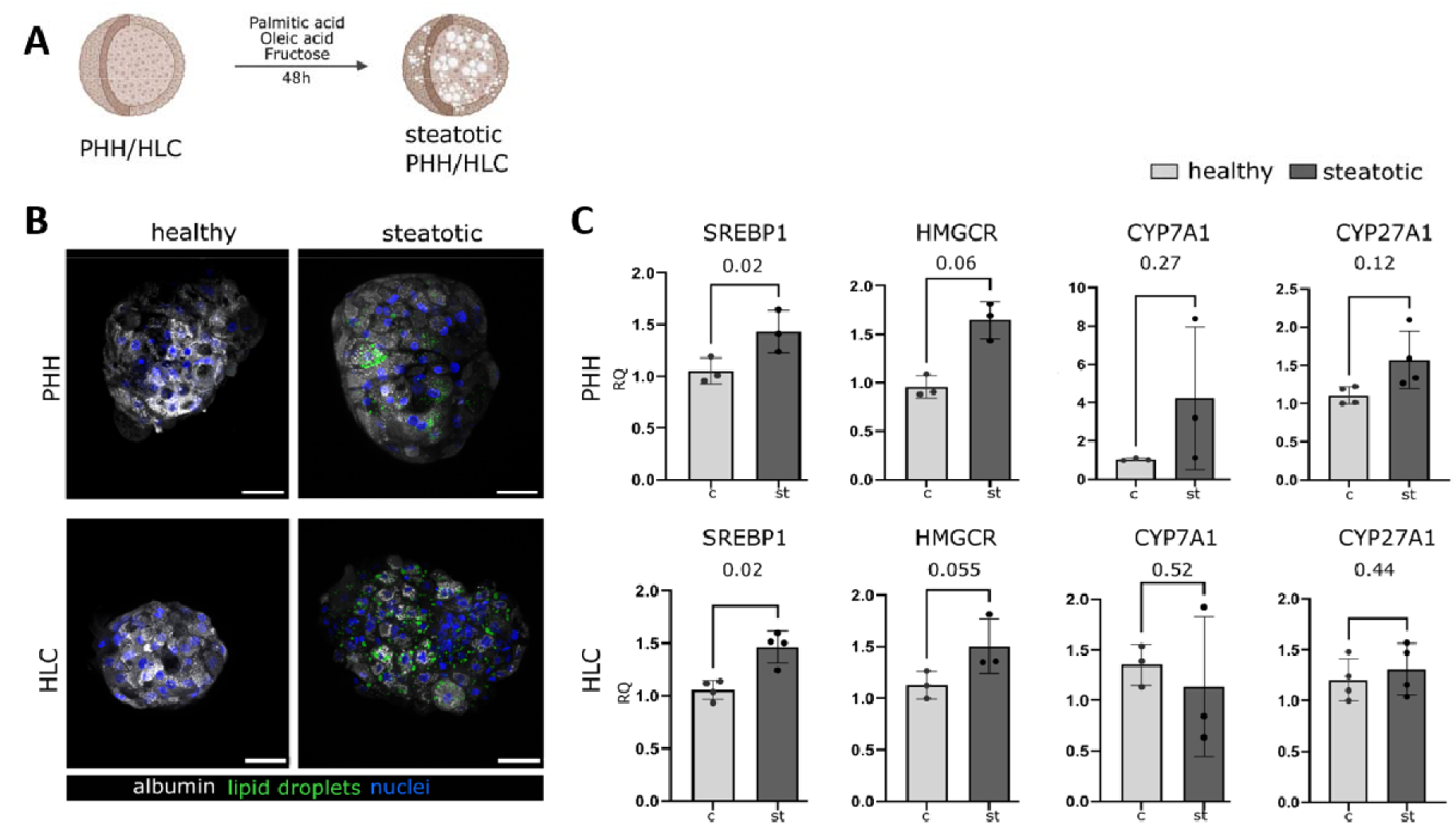
Characterization of hepatic steatosis models using PHH and HLC. A) Schematic presentation of the protocol for inducing hepatic steatosis in vitro. b) Representative images of whole-mount immunofluorescent staining of albumin (white) and cytoplasmic lipids (green) in PHH and HLC. Scale bar 50 μm. c) gene expression in PHH (upper row) and in HLC (low row) after 48 h of steatosis induction. Results represent mean ± SD; n = 3 samples of PHH (N = 2 and more technical replicates for each), and n = 3 cell lines of HLC (N = 3 or more technical replicates for each). A two-tailed, paired t-test is used for the statistics.

### Steatotic liver spheroids/organoids secrete increased levels of oxysterols

To analyse the secretion of oxysterols under pathological conditions, we established a model of hepatic steatosis *in vitro* using spheroids generated from primary human hepatocytes (PHH) and organoids generated from hESC/hiPSC-derived hepatocyte-like cells (HLC) (Figure 7A). The liver spheroids/organoids were characterized by an accumulation of lipids as visualized by Nitrile Red staining, while preserving albumin expression (Figure 7B). Moreover, steatotic PHH and HLC were characterized by a statistically significant upregulation of the expression of sterol regulatory element-binding protein 1 (*SREBP-1*) and a consistent trend of increased expression of HMG-CoA reductase - a rate-limiting enzyme of cholesterol biosynthesis (Figure 7C). We detected no statistically significant difference in the expression of enzymes responsible for oxysterol oxidation (*CYP7A1* and *CYP27A1*) in steatotic PHH and HLC.

Oxysterol levels in media of healthy control spheroids/organoids and steatotic liver spheroids/organoids were subsequently compared. Independent of the spheroids/organoids or experimental repeat, we found a significant difference in the amount of 26-HC secreted from steatotic PHH and HLC compared to controls (*Figure 7A*). While 26-HC was clearly the dominant oxysterol isomer in PHH and HLC, a majority of the steatotic PHH and HLC samples also contained detectable traces of 25-HC and 24S-HC. This suggests sufficient activity of CH25H and CYP46A1 in addition to CYP27A1 in the spheroids/organoids to reach detectable levels of these hydroxycholesterol analytes (Figure 7B). Elevated 26-HC levels detected in steatotic liver spheroids/organoids are in accordance with observations by Ikegami et al. where elevated levels of 26-HC in serum from NAFLD patients [9] was demonstrated. Our findings also support that levels of secreted oxysterols may differ from oxysterols that are retained in cells or liver tissue, if compared to the oxysterols Raselli et al found at elevated levels in liver biopsies from NASH-patients and liver tissue from murine feeding models [10]. In contrast to healthy liver spheroids/organoids, dihydroxycholesterols 7β,25-diHC, 7α,25-diHC, 7α,26-diHC, and 7β,26-diHC were detected in steatotic liver spheroid/organoid samples (*Figure 7C*). In addition, 7α26-HC was present in amounts close to quantification limits in the HLC_2 samples (Figure 8C), demonstrating low but functional expression and activity of CYP7B1. These data indicate a fingerprint of hydroxycholesterols and dihydroxycholesterols that are secreted by the steatotic liver spheroids/organoids and that differentiates them from healthy liver spheroids/organoids.

**Figure 7.**
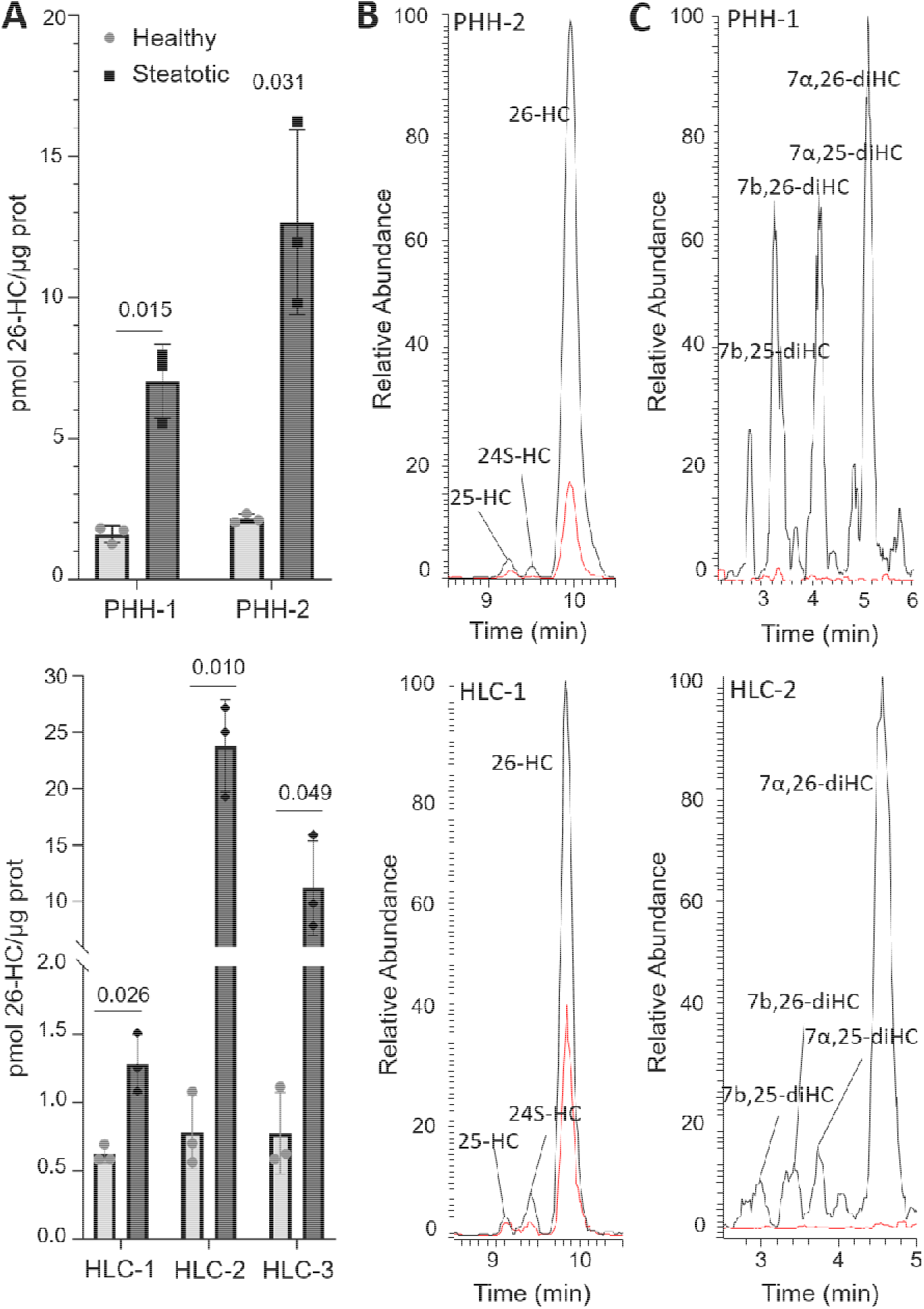
Oxysterol secretion of liver spheroids/organoids. A) The amounts of 26-HC secreted from healthy and steatotic liver spheroids/organoids. The above figure shows two different PHH cell lines, while the below shows three different HLC cell lines. Results represent mean ± SD; n=3 for each cell line. A multiple, unpaired t-test is used to compare healthy and steatotic spheroids/organoids, and P-value is shown for each. B) Representative chromatogram for hydroxycholesterols secreted from healthy (red line) and steatotic (black line) spheroid/organoid samples, PHH (above) and HLC (below), m/z = 514.4 -> 455.4 (hydroxycholesterols). The abundances are normalized to steatotic samples. C) Representative chromatograms for dihydroxycholesterols secreted from healthy control (red line) and steatotic (black line) samples, PHH (above) and HLC (below), m/z = 530.4 -> 471.4. Abundance is normalized to steatotic samples.

**Figure 8.**
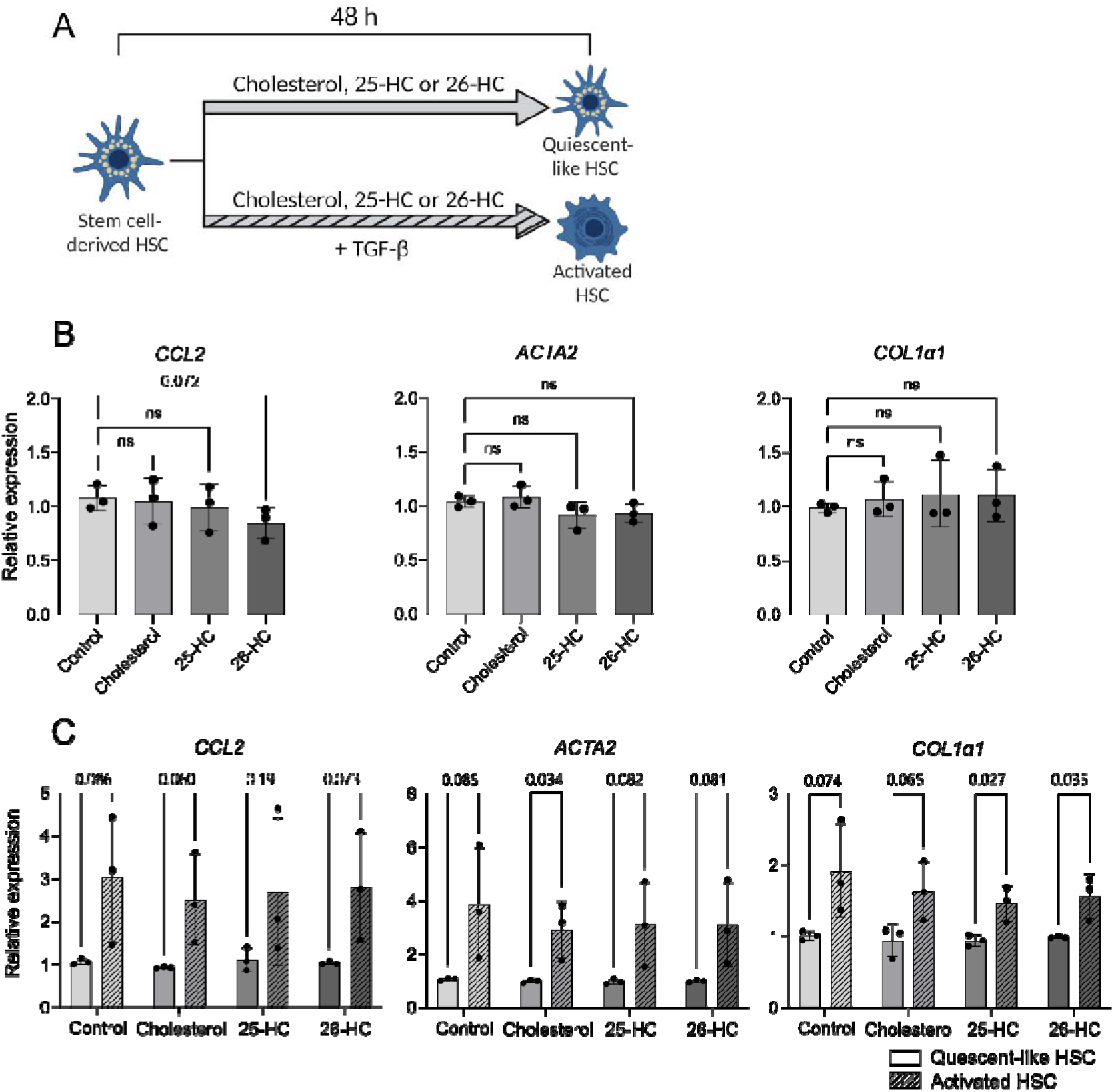
The effect of cholesterol, 25-HC and 26-HC on HSCs. A) Schematic representation of the experiment. HSCs were treated with 50 μM cholesterol, 100 ⊓M 25-HC or 100 ⊓M 26-HC for 48 hours in the presence and absence of activation-inducing stimuli by 25 ng/mL TGF-β. B) Gene expression of activation-associated genes ACTA2, COL1α1 and CCL2 in response to treatment with cholesterol, 25-HC and 26-HC. Gene expression levels were normalized to the control condition. C) Upregulation of activation-associated genes ACTA2, COL1α1 and CCL2 in response to activation-inducing stimuli by TGF-β. Gene expression levels were normalized to “Quiescent-like HSC” samples of each condition. Results represent mean SD, n = 3 independent differentiations of three different PSC lines, three biological replicates for each point. GAPDH was used as the housekeeping gene in all RT-qPCR analyses. Statistical tests: For figure B, a one-way ANOVA with Dunnett’s multiple comparison test comparing control and treatment; for figure C, unpaired, two-tailed t-tests comparing “Quiescent-like HSC” and “Activated HSC” in each group.

### The effect of oxysterols on hepatic stellate cells

The transition between NAFL (steatosis) and NASH (steatohepatitis) is orchestrated by HSCs, which are the main secretors of extracellular matrix and important actors in the recruitment of circulating monocytes [26]. In the homeostatic liver, HSCs exhibit a quiescent phenotype but in response to hepatic damage, including steatosis, HSCs are activated and release extracellular matrix components like collagens as well as pro-inflammatory cytokines such as chemokine (C-C motif) ligand 2 (CCL2) [26–28]. Stem cell-derived HSCs differentiated from one hESC line and two hiPSC lines were used to investigate the impact of cholesterol, 25-HC and 26-HC on the quiescent state of HSCs. 7α,26-diHC, which was strongly released by both steatotic PHH and HLC, was not commercially available and was not evaluated.

Quiescent-like HSCs were treated with cholesterol, 25-HC, or 26-HC (Figure 9A) and mRNA expression detected with RT-qPCR analysis found a consistent trend of *CCL2* downregulation in response to the treatment with 26-HC (Figure 9B). In the same cells, expression of genes associated with the activation of HSCs (*ACTA2*, encoding α-smooth muscle actin, and *COL1α1*, encoding type I collagen) were not altered by cholesterol, 25-HC, or 26-HC (Figure 9B). Moreover, treatment with cholesterol, 25-HC or 26-HC did not appear to prevent or enhance the TGF-β-induced upregulation of *CCL2, ACTA2* and *COL1α1* with the exception of minor variations (Figure 9C).

Collectively, the results suggest that 26-HC may exert an anti-inflammatory signal on quiescent HSCs and consequently might have an impact on the progression of NAFL to NASH, as evident by consistent trends in lowered expression of *CCL2*. However, the effect was absent in TGF-β-activated HSCs. These observations are consistent with existing literature that implicates 26-HC in the attenuation of NAFL [29].

## CONCLUSION

As a new analytical approach, we have introduced sterolomics to organoids, an emerging tool for e.g. replacing animal models in biomedical research. Our targeted LC-MS method for determination of Girard T derivatized oxysterols reveal that organoids secrete oxysterols, and 26-HC and diHCs were significantly elevated in steatotic samples. Our results, using human liver organoid models, support previous findings based on human serum samples, adding arguments to the validity of human liver organoids as disease models. The association of specific oxysterols with a potential anti-inflammatory role calls for further investigations of oxysterols in NAFLD disease progression. As the elevated oxysterol levels do not correspond with mRNA findings in steatotic spheroids/organoids it shows the importance of direct measurements of metabolites.

## ACKNOWLEDGEMENTS

This work was supported by the Research Council of Norway through its Centre of Excellence scheme, project number 262613, and project number: 295910 (National Network of Advanced Proteomics Infrastructure). Support was also provided by UiO:Life Science and the Olav Thon Foundation.

## Notes

### Competing Interest Statement

The authors have declared no competing interest.

### Summary of Updates

Additional ORCID information added to author list.

## References

1. Moore, J.B., From sugar to liver fat and public health: systems biology driven studies in understanding non-alcoholic fatty liver disease pathogenesis. Proceedings of the Nutrition Society, 2019. 78(3): p. 290–304.

2. Younossi, Z.M., Non-alcoholic fatty liver disease–a global public health perspective. Journal of hepatology, 2019. 70(3): p. 531–544.

3. Le, M.H., et al., 2019 global NAFLD prevalence-A systematic review and meta-analysis. Clinical Gastroenterology and Hepatology, 2021.

4. Stefan, N., H.-U. Häring, and K. Cusi, Non-alcoholic fatty liver disease: causes, diagnosis, cardiometabolic consequences, and treatment strategies. The Lancet Diabetes & Endocrinology, 2019. 7(4): p. 313–324.

5. Gallage, S., et al., A researcher’s guide to preclinical mouse NASH models. Nature Metabolism, 2022: p. 1–18.

6. Method of the Year 2017: Organoids. Nature Methods, 2018. 15(1): p. 1–1.

7. Kozyra, M., et al., Human hepatic 3D spheroids as a model for steatosis and insulin resistance. Scientific reports, 2018. 8(1): p. 1–12.

8. Jia, W., et al., Targeting the alternative bile acid synthetic pathway for metabolic diseases. Protein & cell, 2021. 12(5): p. 411–425.

9. Ikegami, T., et al., Increased serum liver X receptor ligand oxysterols in patients with non-alcoholic fatty liver disease. Journal of gastroenterology, 2012. 47.

10. Raselli, T., et al., Elevated oxysterol levels in human and mouse livers reflect nonalcoholic steatohepatitis[S]. Journal of Lipid Research, 2019. 60(7): p. 1270–1283.

11. Griffiths, W.J. and Y. Wang, Oxysterol research: a brief review. Biochemical Society Transactions, 2019. 47(2): p. 517–526.

12. Lehmann, J.M., et al., Activation of the nuclear receptor LXR by oxysterols defines a new hormone response pathway. Journal of Biological Chemistry, 1997. 272(6): p. 3137–3140.

13. Becares, N., et al., Impaired LXRa Phosphorylation Attenuates Progression of Fatty Liver Disease. Cell Reports, 2019. 26(4): p. 984–995.e6.

14. Pandak, W.M. and G. Kakiyama, The acidic pathway of bile acid synthesis: Not just an alternative pathway. Liver research, 2019. 3(2): p. 88–98.

15. Fakheri, R.J. and N.B. Javitt, 27-Hydroxycholesterol, does it exist? On the nomenclature and stereochemistry of 26-hydroxylated sterols. Steroids, 2012. 77(6): p. 575–577.

16. Busek, M., et al., Pump-less, recirculating organ-on-a-chip (rOoC) platform. Lab on a Chip, 2023.

17. Coll, M., et al., Generation of hepatic stellate cells from human pluripotent stem cells enables in vitro modeling of liver fibrosis. Cell stem cell, 2018. 23(1): p. 101–113. e7.

18. Vallverdú, J., et al., Directed differentiation of human induced pluripotent stem cells to hepatic stellate cells. Nature Protocols, 2021.16(5): p. 2542–2563.

19. Schindelin, J., et al., Fiji: an open-source platform for biological-image analysis. Nature methods, 2012. 9(7): p. 676–682.

20. Roberg-Larsen, H., et al., Highly automated nano-LC/MS-based approach for thousand cell-scale quantification of side chain-hydroxylated oxysterols [S]. Journal of lipid research, 2014. 55(7): p. 1531–1536.

21. Griffiths, W.J., et al., Discovering oxysterols in plasma: a window on the metabolome. Journal of proteome research, 2008. 7(8): p. 3602–3612.

22. Roberg-Larsen, H., et al., Highly automated nano-LC/MS-based approach for thousand cell-scale quantification of side chain-hydroxylated oxysterols. J Lipid Res, 2014. 55(7): p. 1531–6.

23. Solheim, S., et al., Fast liquid chromatography-mass spectrometry reveals side chain oxysterol heterogeneity in breast cancer tumour samples. The Journal of Steroid Biochemistry and Molecular Biology, 2019. 192: p. 105309.

24. Roberg-Larsen, H., et al., High sensitivity measurements of active oxysterols with automated filtration/filter backflush-solid phase extraction-liquid chromatography–mass spectrometry. Journal of Chromatography A, 2012. 1255: p. 291–297.

25. Graffmann, N., B. Scherer, and J. Adjaye, In vitro differentiation of pluripotent stem cells into hepatocyte like cells-basic principles and current progress. Stem Cell Research, 2022: p. 102763.

26. Tsuchida, T. and S.L. Friedman, Mechanisms of hepatic stellate cell activation. Nature reviews Gastroenterology & hepatology, 2017. 14(7): p. 397–411.

27. Kitto, L.J. and N.C. Henderson, Hepatic stellate cell regulation of liver regeneration and repair. Hepatology Communications, 2021. 5(3): p. 358–370.

28. Tacke, F., Targeting hepatic macrophages to treat liver diseases. Journal of hepatology, 2017. 66(6): p. 1300–1312.

29. Bieghs, V., et al., The cholesterol derivative 27-hydroxycholesterol reduces steatohepatitis in mice. Gastroenterology, 2013. 144(1): p. 167–178. e1.

