## Supporting Information for "Mass Spectrometry Reveals that Oxysterols are Secreted from Non-Alcoholic Fatty Liver Disease Induced Organoids"

**MS parameters**

The TSQ Vantage triple quadrupole mass spectrometer was operated in positive mode, with a voltage of +300 V, collision gas pressure of 1.0 mTorr, a sheat gas pressure of 60 and aux valve flow of 10. Capillary temperature was 380 °C and vaporizer temperature was 300 °C. All analytes were monitored in selected reaction mode (SRM), see Table 1for more information.

Table 1 SRM-transitions and collision energy for all targeted analytes. For collision energy the two listed values correlates to the quantifier m/z and qualifier m/z, respectively.

| Analytes | Parent m/z | Collision energy | Quantifier m/z | Qualifier m/z |
| --- | --- | --- | --- | --- |
| 22R-HC  24S-HC  25-HC  26-HC | 514.4 | 36/30 | 455.4 | 427.4 |
| 7α,25-diHC  7α,26-diHC  7β,26-diHC  7β,25-diHC  7α,24S-diHC | 530.4 | 35/32 | 471.4 | 443.4 |
| 25-HC-d_6_  27-HC-d_6_ | 521.4 | 30 | 462.4 | 434.4 |
| 7α,25-diHC-d_6_  7α,26-diHC-d_6_ | 536.4 | 35/32 | 477.4 | 449.4 |
| Cholesterol autoxidation monitoring | 517.4 | 30 | 458.4 |  |

**Method evaluation**

Table 2 The data for evaluation of method, showing calibration curve-equation with R^2^-values, and the calculated precision for LOQs both for intra-day and inter-day and for three different concentrations. Intra-day precision is calculated using one-way Anova, and is based on three replicates at four different days.

| Analyte | Range | Y=ax+b (R^2^) | Precision LLOQ  (50 pM) | | Precision MLOQ  (200 pM) | | Precision HLQ  (500 pM) | |
| --- | --- | --- | --- | --- | --- | --- | --- | --- |
|  | pM |  | Intra-day (n=6) | Inter-day (n=4) | Intra-day (n=6) | Inter-day (n=4) | Intra-day (n=6) | Inter-day (n=4) |
| 22R-HC | 50-500 | 0.9899 | 44 | 15 | 12 | 8 | 10 | 10 |
| 25-HC | 50-500 | 0.9979 | 10 | 13 | 8 | 3 | 7 | 6 |
| 24S-HC | 50-500 | 0.9899 | 17 | 18 | 3 | 8 | 6 | 6 |
| 26-HC | 50-500 | 0.9985 | 13 | 14 | 13 | 12 | 7 | 6 |
| 7α,25-diHC | 50-500 | 0.9879 | 16 | x | 8 | 26 | 28 | 27 |
| 7α,26-diHC | 50-500 | 0.9977 | 19 | 48 | 17 | 31 | 15 | 36 |
| 7β,26-diHC | 50-500 | 0.9964 | 22 | 38 | 8 | 25 | 13 | 27 |
| 7β,25-diHC | 50-500 | 0.9742 | 17 | 15 | 7 | 8 | 23 | 20 |
| 7α,24S-diHC | 50-500 | 0.9759 | 7 | 17 | 14 | 22 | 21 | 41 |

**sc-HSC characterization**

(A) Schematic representation of the sc-HSC differentiation protocol.
(B) Representative immunofluorescence images of known HSC-markers PDGFR-β, Vimentin and NCAM1 in sc-HSCs. Scale bars: 40 µm.
(C) Representative bright field- and UV images of sc-HSCs, indirectly imaging the storage of vitamin A.

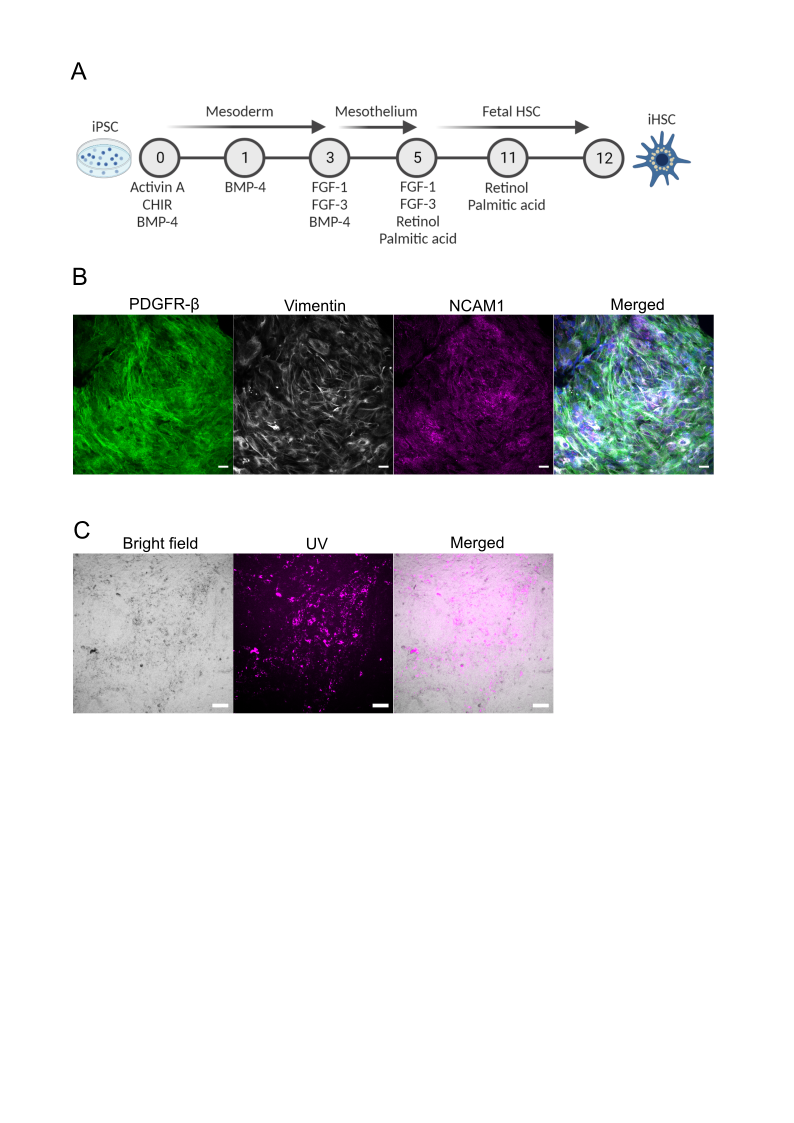
